# SKIN AS A POTENTIAL ENTRY POINT FOR SARS-COV-2

**DOI:** 10.64898/2026.04.07.717019

**Authors:** Dmitry Trubetskoy, Patrick Grudzien, Daria A. Chudakova, Anna Klopot, Bo Shi, Pankaj Bhalla, Bethany Perez White, Irina Budunova

## Abstract

The primary route of SARS-CoV-2 entry is via respiratory epithelium. However, many COVID-19 patients developed dermatological lesions, and SARS-CoV-2 RNA has been detected in the patients’ skin. Inflammatory skin diseases, psoriasis and atopic dermatitis (AD), significantly increased the risk of COVID-19. To evaluate the potential role of skin in SARS-CoV-2 host interactions, we utilized 3D human skin organoids (HSO) generated from human epidermal keratinocytes, as well as neonatal skin explants. HSO were treated with cytokines involved in acute and chronic skin inflammation and cytokine storm in severe COVID-19 disease, TNF-α, IL-6, IL-1β, and IFN-γ, individually and in combination. HSO were also treated with Th1 (TNF-α + IL-17) and Th2 (IL-4 + IL-13) cocktails inducing pro-psoriasis and pro-AD HSO changes, respectively. All individual cytokines, and especially their combinations, elevated the expression of ACE2 and TMPRSS2 at mRNA/protein levels. The Th2 induced only TMPRSS2, the Th1 predominantly induced ACE2. Topically applied Spike-pseudotyped lentiviral Tomato reporter, which binds ACE2 similarly to SARS-CoV-2, successfully infected control and cytokine-treated HSO as well as neonatal skin explants. Cytokine treatment, especially TNF-α + IL-6 + IL-1β + IFN-γ and the Th1, significantly increased viral entry. Transcriptomic analysis further revealed partial overlap between gene expression signatures induced by Spike-mediated entry in inflamed HSO and those observed in lung tissue from COVID-19 patients, supporting the biological relevance of skin models. Together, these findings demonstrate that inflammation enhances the permissiveness of human skin to SARS-CoV-2 entry, suggesting that the skin may represent a previously underappreciated interface in viral host interactions.

## 1. Introduction

The Coronavirus Disease 2019 (COVID-19), caused by Severe acute respiratory syndrome coronavirus 2 (SARS-CoV-2), has posed a major global health challenge and continues to represent a potential threat due to the ongoing emergence of SARS-CoV-2 variants with differing levels of immune evasion and pathogenicity. Despite extensive research efforts, many important aspects of SARS-CoV-2 biology remain incompletely understood, including potential alternative routes of transmission and the mechanisms underlying the diverse clinical manifestations of COVID-19.

SARS-CoV-2 is a single-stranded RNA virus whose infectivity mainly depends on the binding of the viral Spike (S) glycoprotein to the host cell receptor angiotensin-converting enzyme 2 (ACE2), followed by priming and cleavage by the host protease transmembrane serine protease 2 (TMPRSS2) [1,2].

ACE2, a monocarboxypeptidase involved in renin-angiotensin signaling, plays important roles in cardiovascular and renal physiology. TMPRSS2 is a membrane-associated protease with major functions in prostate biology and prostate cancer progression. TMPRSS2 promotes SARS-CoV-2 infection by cleaving ACE2, thereby enhancing viral uptake, and by cleaving the Spike protein itself, enabling membrane fusion and viral internalization [1,2]. There are also several other non-primary receptors and cofactors that may contribute to SARS-CoV-2 cell entry [3].

The respiratory epithelium is recognized as the primary entry point for SARS-CoV-2. However, cells in other tissues and organs including the gastrointestinal tract, kidney, heart, eye, and skin also express ACE2 and TMPRSS2 [4-9]. This broad expression pattern may underline the virus’s multi-organ tropism and contribute to the extra-pulmonary clinical manifestations observed in COVID-19 patients.

The role of barrier organs such as the gastrointestinal tract and eye as additional entry points for SARS-CoV-2 has been established [5,10], but the potential role of skin remains insufficiently investigated. Recent single-cell RNA sequencing and immunostaining studies demonstrated that ACE2 is expressed in both basal and differentiated keratinocytes and is co-expressed with TMPRSS2 in the stratum granulosum [8, 11]. According to our bulk RNA-sequencing (RNA-seq) data, both ACE2 and TMPRSS2 rank within the top 10% of expressed genes in healthy human skin [12]. Moreover, the levels of ACE2 mRNA in skin are somewhat comparable with its levels in lungs, the primary entry site of SARS-CoV-2 [13], highlighting skin as a potentially permissive tissue for SARS-CoV-2 infection.

Despite its efficient physical and immunological barrier properties, skin is directly infected by viruses such as Herpes simplex virus, Varicella zoster virus, and Molluscum contagiosum virus, whose entry and replication largely occur within the epidermis. There is also evidence suggesting that skin may be targeted by SARS-CoV-2. Indeed, 1–20% of COVID-19 patients develop transient or persistent skin manifestations, including chilblain-like acral lesions (pernio, also known as “COVID toes”), vasculitis-like lesions, urticaria-like lesions, and maculopapular eruptions, which can occur even before the onset of major systemic symptoms [14-16]. Further, viral RNA and capsid proteins of SARS-CoV-2 have been detected in skin biopsies and autopsy samples from COVID-19 patients [15], [17-18].

Importantly, inflammatory skin diseases such as psoriasis and atopic dermatitis (AD) have been associated with an increased risk of COVID-19 [19-21]. Moreover, increased ACE2 and TMPRSS2 expression has been reported in lesional skin of psoriatic patients [20, 22-23]. However, the pattern and dynamic of the changes of the ACE2 and TMPRSS2 expression under different inflammatory conditions in skin, and the role of healthy or inflamed human skin in SARS-CoV-2 entry and COVID-19 pathogenesis remains poorly understood.

The major goals of our study were to: (i) determine whether SARS-CoV-2 can infect skin; and (ii) assess how individual inflammatory cytokines and cytokine cocktails: Th1 (TNF-α+IL17), Th2 (IL4+IL13), and a combination of IL-1β + IL-6 + IFN-γ + TNF-α resembling the inflammatory cytokine storm induced by SARS-CoV-2 infection, affect ACE2 and TMPRSS2 expression and infection efficiency. We employed two in vitro skin models: neonatal skin explants and human skin organoids (HSO) generated from neonatal keratinocytes seeded on a collagen matrix and maintained on liquid/air interface [24]. Mature HSO have well established multilayer epidermis closely resembling epidermis of human skin [24]. To visualize and quantify infection, we used a Spike-pseudotyped lentiviral Tomato reporter expressing the Spike protein from the original Wuhan-Hu-1 strain.

Given the reported higher risk of COVID-19 infection in African American (AA) populations [25-27], and different risk of inflammatory skin diseases such as AD and psoriasis in AA and White non-Hispanic (WNH) populations we also evaluated potential differences in ACE2 and TMPRSS2 induction by cytokines and infection rates in AA and WNH skin models.

## 2. Results

### 2.1. ACE2 and TMPRSS2 Are Highly Expressed in Human Skin and 3D Human Skin Organoids

Our previously published transcriptome analysis of healthy human skin revealed robust expression of both ACE2 and TMPRSS2 at the mRNA level, with both genes ranking among the top 10% of expressed genes in human skin [12,28]. Here, we extended these findings by direct analysis of ACE2 and TMPRSS2 expression by qRT-PCR and Western blotting in neonatal skin and in 3D human skin organoids (HSO) that develop all epidermal layers including basal, spinous, granular layers and extensive stratum corneum critical for skin barrier function (Fig. 1A).

**Figure 1.**
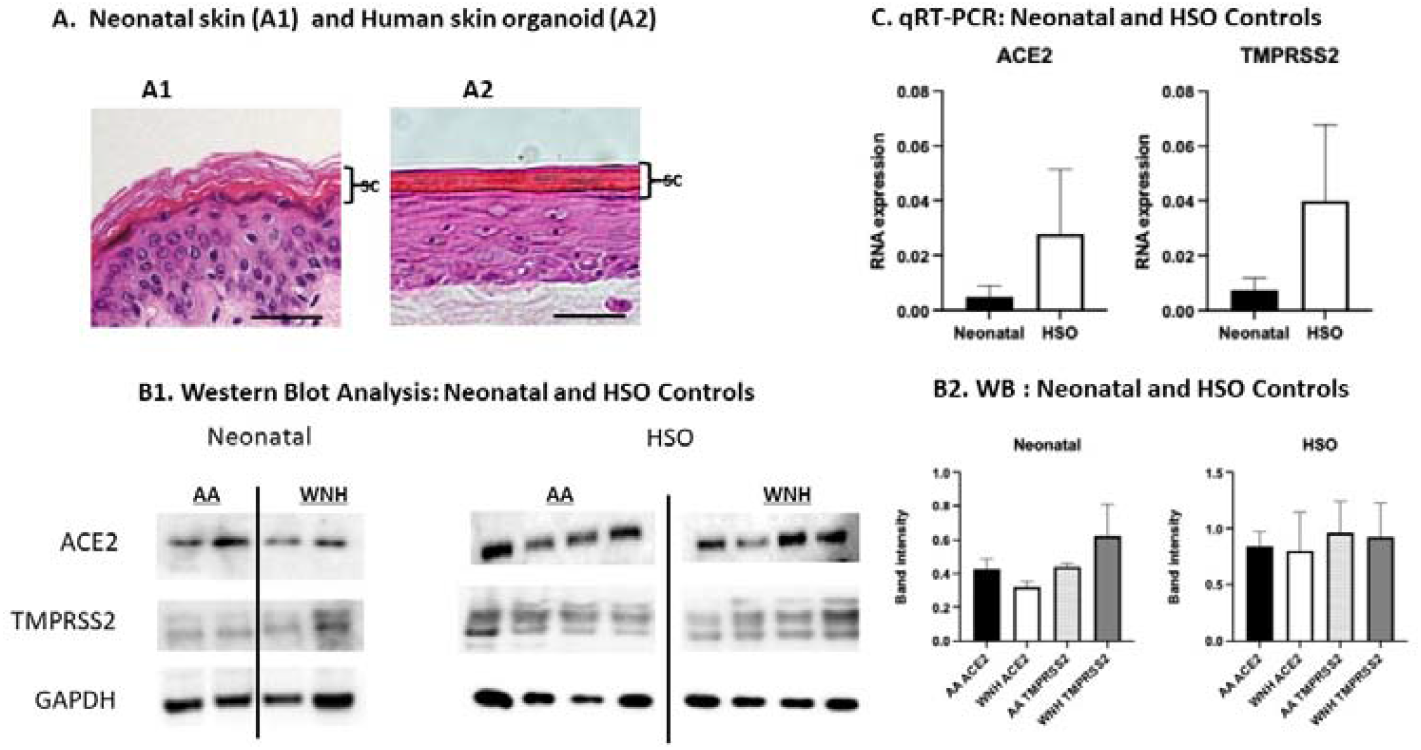
Expression of ACE2 and TMPRSS2 in human neonatal skin and in HSO. RNA and proteins were extracted from untreated AA and WNH neonatal foreskin and HSO epidermis. (A). Morphology of epidermis in neonatal skin and HSO. H&E staining, Scale Bar is 10 μm. SC -stratum corneum (B). ACE2 and TMPRSS2 expression analyzed by Western blotting in individual samples. GAPDH was used as a loading control; (C). qRT-PCR analysis of *ACE2* and *TMPRSS2* expression. *RPL27* was used as a cDNA normalization control. Statistical analysis was done by an unpaired two-tailed Welch’s t-test. Error bars represent mean ± SD.

These experiments also revealed big inter-individual variability in ACE2 and TMPRSS2 expression (Fig. 1B and Fig.2C). We did not observe significant differences in ACE2 or TMPRSS2 expression between control AA and WNH skin explants or between control AA and WNH HSO due to a small number of samples (Fig 1B), However, the experiments with a larger number of samples, revealed a higher statistically significant expression of TMPRSS2 in AA HSO (Fig. 2C, control).

**Figure 2.**
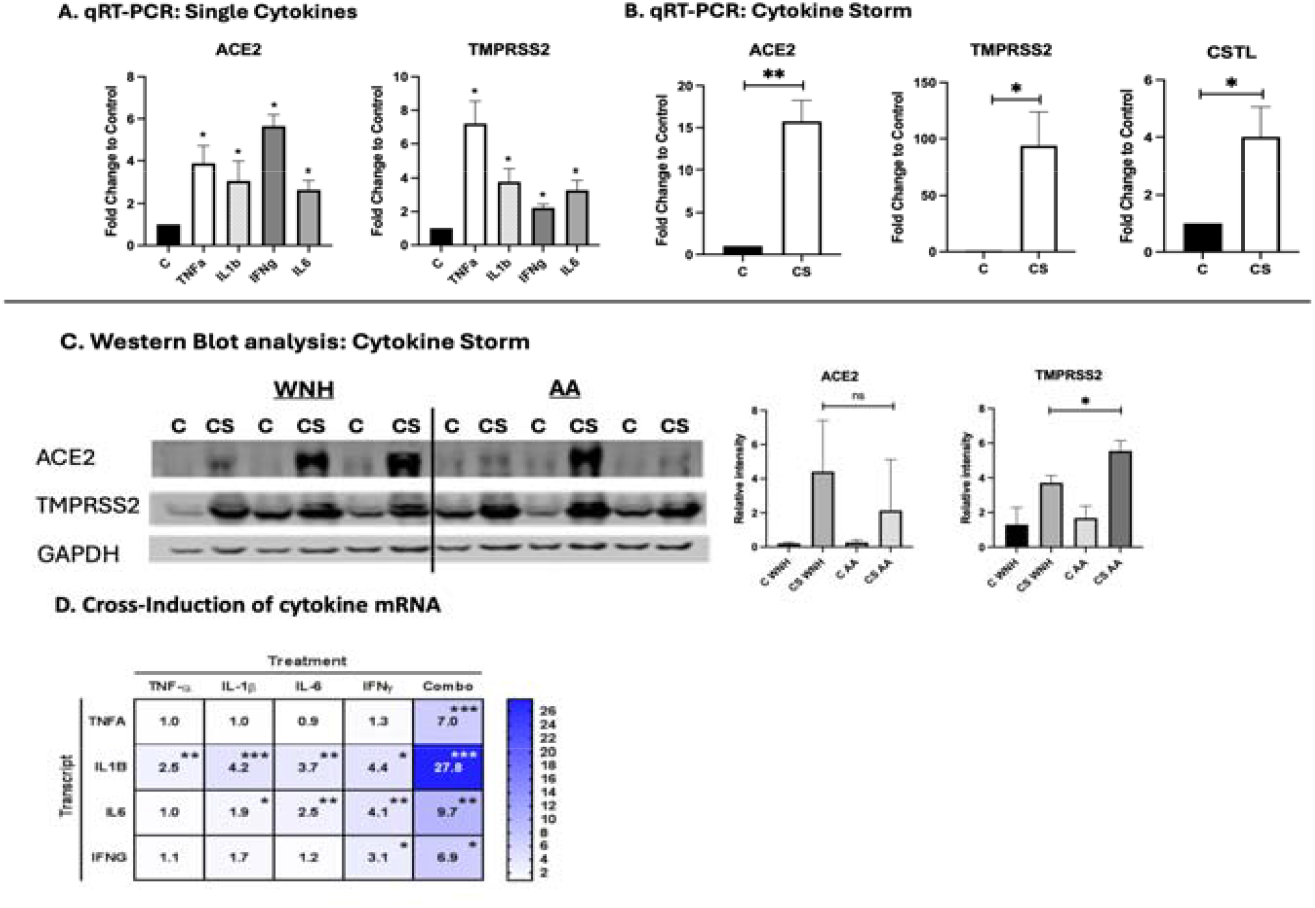
Induction of ACE2 and TMPRSS2 in HSO by cytokines related to the COVID-19 cytokine storm. Mature HSO were treated with TNF-α, IL-6, IL-1β, and IFN-γ individually (10 ng/mL each) or in combination for 7 days. (A, B) ACE2 and TMPRSS2 expression was analyzed by qRT-PCR. RPL27 used as a cDNA normalization control; (C) Western blot analysis of ACE2 and TMPRSS2 expression. GAPDH was used as a loading control, (D) HSO (3-4 HSO/group made from the individual keratinocyte cell lines) were treated with the cytokine storm - related individual cytokines as in A. The cross-induction of TNF-α, IL-6, IL-1β, and IFN-γ expression was assessed by qRT-PCR. The results are presented as Heatmap. Color intensity reflects fold change in mRNA expression induction compared to internal control. C-Control, CS - cytokine storm-related cytokine combination. Statistical analysis of was performed by an unpaired two-tailed Welch’s t-test. Error bars represent mean ± SD. Statistically significant difference compared to control: *-P<0.05; ** - P<0.02; *** - P<0.01; ns – not significant.

### 2.2. Proinflammatory Cytokines Strongly Induce ACE2 and TMPRSS2 Expression in HSO

To determine whether inflammation affects key factors central to SARS-CoV-2 entry, mature HSO cultures were treated with TNF-α, IL-6, IL-1β, and IFN-γ cytokines that play key roles in acute and chronic skin inflammation and in the cytokine storm associated with severe COVID-19 [29-30]. All four cytokines significantly upregulated ACE2 and TMPRSS2 expression at both the mRNA and protein levels, with particularly strong induction when applied in combination (Fig. 2A, B, C). The synergistic effects of cytokine combination on ACE2 and TMPRSS2 expression reflect positive crosstalk among these cytokines, as each cytokine upregulated the expression of the others (Fig. 2D).

The magnitude of induction varied significantly among individual HSO, depending on the donor source of the primary keratinocytes. The induction of ACE2 at the protein level was comparable in AA and WNH HSO, but TMPRSS2 induction was higher in AA HSO (Fig. 2).

Next, we examined the effect of cytokine treatment on the expression of other reported receptors for the SARS-CoV-2 Spike protein and alternative proteases implicated in viral entry, including Kringle Containing Transmembrane Protein 1 (KREMEN1), Asialoglycoprotein Receptor 1 (ASGR1), AXL receptor tyrosine kinase, cathepsin L (CTSL), furin, and neuropilin-1 (NPL1) [31]. This analysis was performed using our previously generated RNA-seq data from HSO treated with TNF-α, IL-6, IL-1β, and IFN-γ [28] (GSE163711, PRJNA1068171). We observed minimal or no changes in the expression of these genes following cytokine treatment, with the exception of CTSL, whose expression was increased by ∼ 5 folds in HSO treated with combination of cytokines (Fig. 2B).

Th1 (TNF-α + IL-17) and Th2 (IL-4 + IL-13) cytokine cocktails are known to induce in HSO cultures pro-psoriasis and pro–atopic dermatitis (AD) morphological and molecular changes, respectively [24],[32]. Because inflammatory skin diseases such as psoriasis and AD were associated with increased COVID-19 risk [21],[33], and because disparities in the prevalence of inflammatory skin diseases have been reported between AA/Black and WNH populations [26], we assessed the effects of Th1 (pro-psoriasis) and Th2 (pro-AD) cytokine cocktails on ACE2 and TMPRSS2 protein expression in AA and WNH HSO groups (Fig. 3).

**Figure 3.**
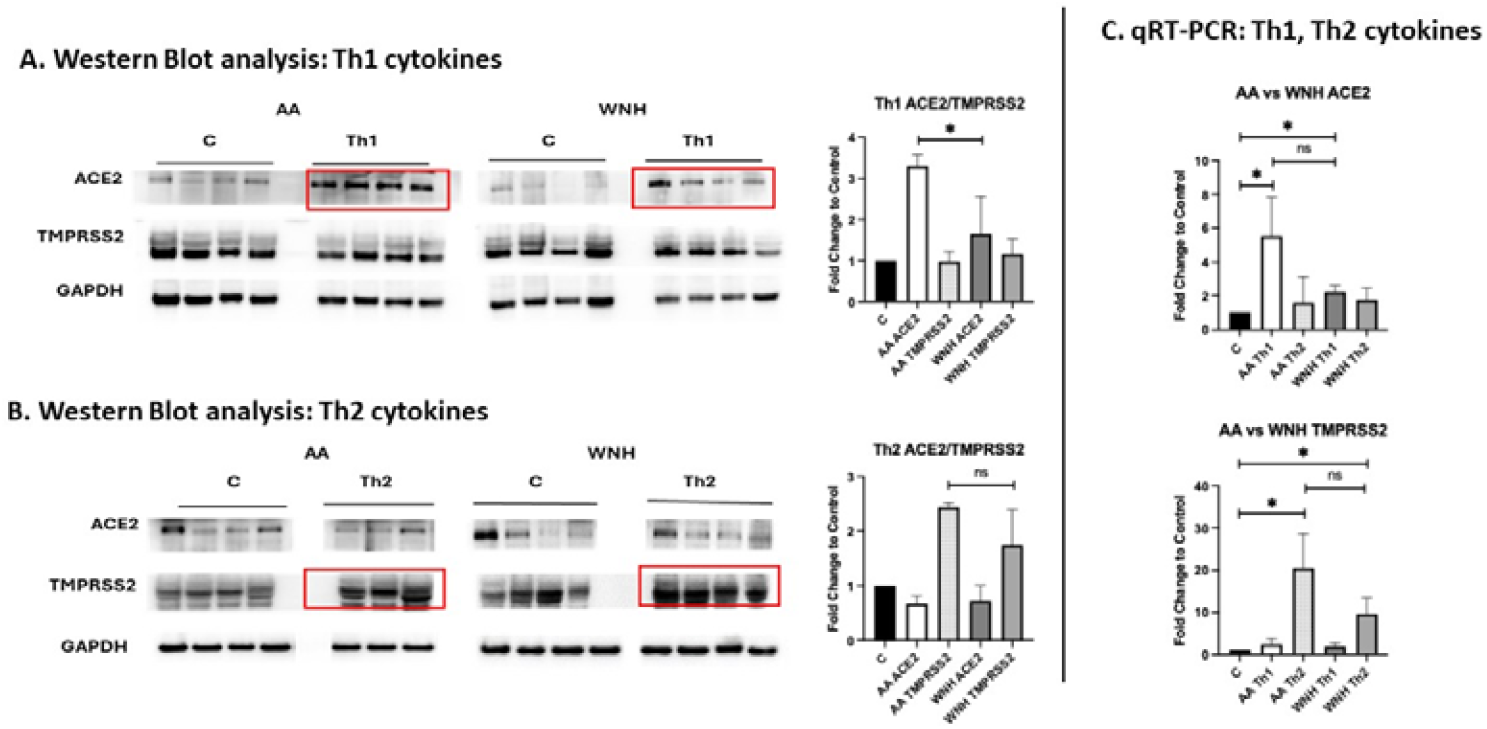
Induction of ACE2 and TMPRSS2 in HSO by Th1 and Th2 cytokine cocktails. Mature AA and WNH HSO were treated with Th1 (IL-17 + TNF-α, 10 ng/mL each) or Th2 (IL-4 + IL-13, 30 ng/mL each) cytokine cocktails for 7 days. (A, B) Western blot analysis of ACE2 and TMPRSS2 expression in AA and WNH HSO after the treatment with Th1 (A) and Th2 (B) cytokines. GAPDH was used as a as loading control; (C) qRT-PCR analysis of ACE2 and TMPRSS2 after Th1 and Th2 treatments, Statistical analysis by an unpaired two-tailed Welch’s t-test. Error bars represent mean ± SD. Statistically significant difference compared to control (P<0.05).

The results of Western blots correlated with induction of ACE2 and TMPRSS2 expression at mRNA level. Interestingly, Th1 cytokines predominantly induced ACE2 expression, whereas Th2 cytokines primarily induced TMPRSS2 expression in HSO. Moreover, the effects of Th1 cytokine cocktail on ACE2 expression at protein level and as a trend at mRNA level were more pronounced in AA HSO (Fig. 3A, B).

### 2.3. Infection of Control and Inflamed Skin Models with Spike-Pseudotyped Tomato/RFP Lentiviral Reporter

The expression of ACE2 and TMPRSS2 in the neonatal skin and in HSO suggested that these skin models could be susceptible to SARS-CoV-2 infection. To test this, we employed a biosafety level 2, non-replicating lentiviral reporter pseudotyped with the SARS-CoV-2 Spike glycoprotein and expressing a Tomato/RFP fluorescent reporter, replacing the commonly used VSV-G envelope protein that binds cells via non-specific lipids. This approach limits viral entry to ACE2-expressing target cells and has been widely used for screening of antiviral compounds and studying host cell infection mechanisms [34].

Topical application of the Spike-pseudotyped Tomato/RFP reporter to mature HSO cultures resulted in strong Tomato/RFP fluorescence within the epidermis 48–72 hours post-infection (Fig. 4A). Similar infection patterns were observed in neonatal skin explants following topical infection with the Spike-pseudotyped Tomato/RFP reporter (Fig. 4A).

**Figure 4.**
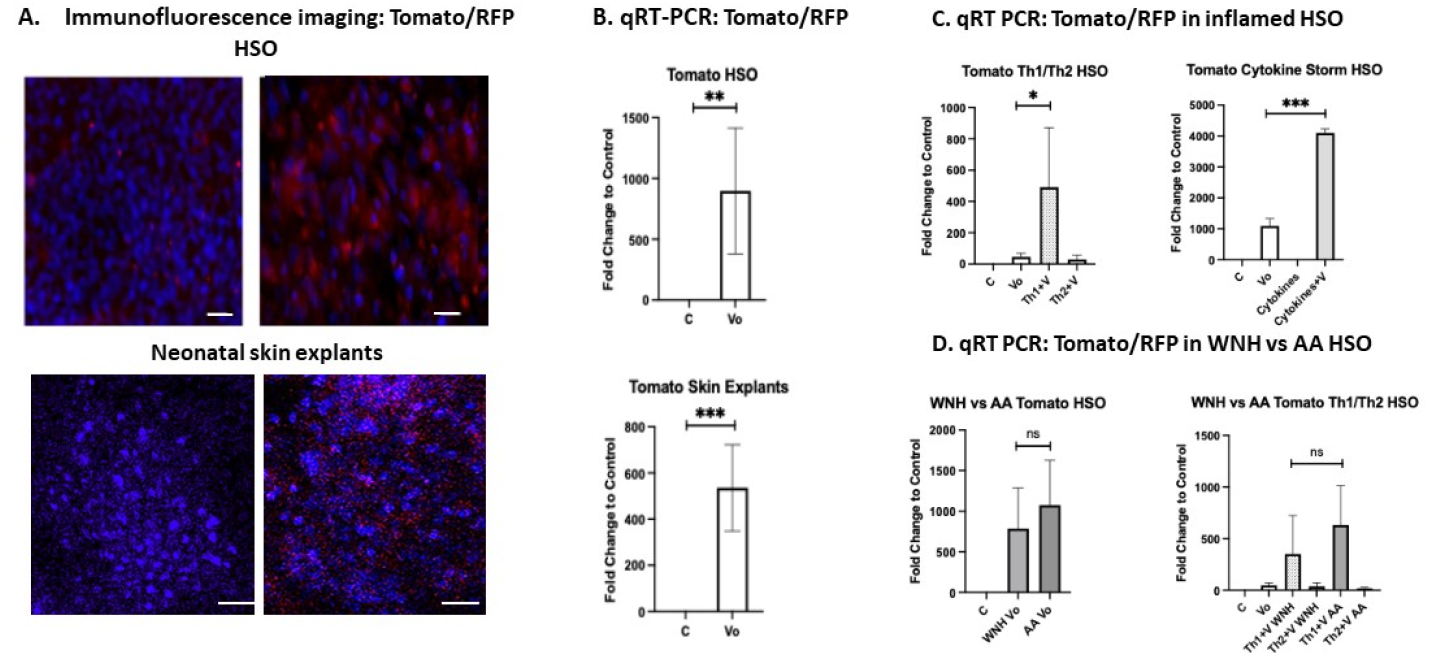
Neonatal skin explants and mature HSO were pretreated with vehicle control, cytokine storm-related cytokines (TNF-α + IL-6 + IL-1β + IFN-γ), Th1, or Th2 cytokines for 3 days. Tomato-expressing lentivirus pseudotyped with the SARS-CoV-2 Spike protein (10 uL, 10^7^ TU /mL) was applied to the surface of the skin models. (A). Tomato/RFP fluorescence was detected 72 hours post-infection in HSO and neonatal skin explants pretreated with vehicle control (Control) or virus only (Vo). Images were acquired from the apical aspect of whole-mount skin explants or HSO epidermis separated from the collagen matrix; (B). Quantification of infection by qRT-PCR analysis of Tomato/RFP mRNA expression in control (C) and virus only (Vo) HSO samples and Skin Explants. (C). Quantification of infection by qRT-PCR analysis of Tomato/RFP mRNA expression in “inflamed” HSO pretreated with cytokines before topical application of pseudotyped lentivirus. The groups: control (C), virus only (Vo), Th1 + virus (Th1+V), Th2 + virus (Th2+V), cytokine storm (cytokines) and cytokine storm + virus (cytokines +Vo). RPL27 was used as a normalization control. Statistical analysis was performed using an unpaired two-tailed Welch’s t-test. Error bars represent mean ± SD. Statistically significant difference compared to control: * - P<0.05; ** - P<0.02; ***P<0.01.

For quantitative analysis of infection efficiency in control and inflamed skin models, we measured Tomato/RFP mRNA expression by qRT-PCR 72 hours after topical viral application (Fig. 4B). Infection rates were dramatically increased in HSO pretreated with the combination of TNF-α + IL-6 + IL-1β + IFN-γ that mimics the cytokine storm observed in severe COVID-19 patients. Th1 cytokine pretreatment also significantly increased infection efficiency, whereas Th2 cytokines did not (Fig. 4C). This was consistent with the preferential induction of ACE2 rather than TMPRSS2 by Th1 cytokines in these models (Fig. 3C). No significant differences in infection rates were observed between AA and WNH inflamed HSO, possibly due to the limited number of individual skin models used in these experiments (Fig. 4D).

### 2.4. Similarity Between Molecular Signatures of inflamed HSO infected by SARS-CoV-2 Reporter and SARS-CoV-2 Infection in Lungs of COVID-19 Patients

To further assess the clinical relevance of our skin model for SARS-CoV2 infection, we compared gene expression signatures from lungs of deceased COVID-19 patients (GSE150316 presented at https://maayanlab.cloud/Enrichr/ site generated by Dr. Ma’ayan’s Laboratory at Mount Sinai Center for Bioinformatics (NYC), with transcriptomic profile of HSO treated by the cytokine storm–related cytokine combination typical for patients with severe COVID-19 and subsequently infected by the SARS-CoV-2 Spike reporter [29-30](Fig. 5).

**Figure 5.**
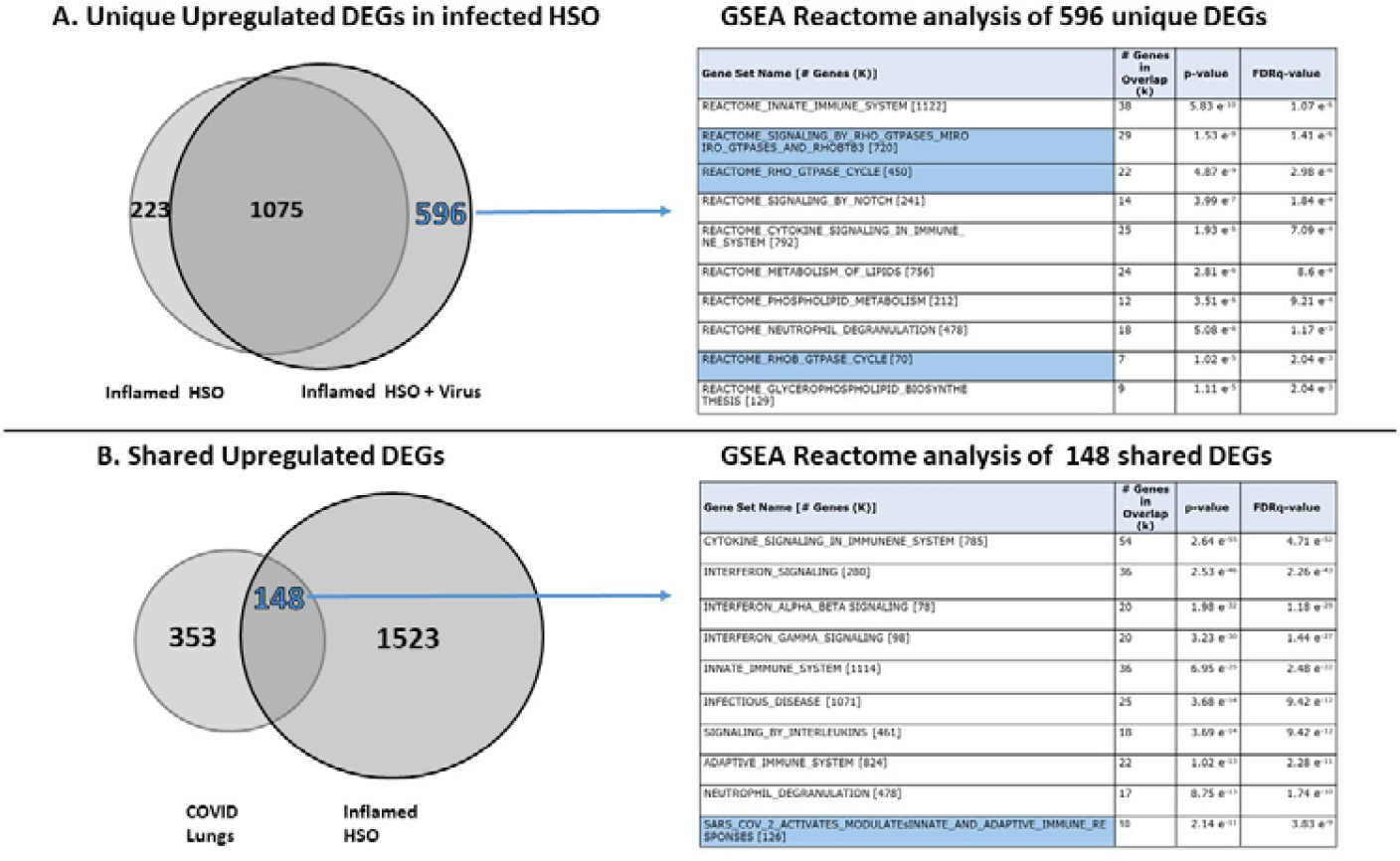
Similarity Between Molecular Signatures of inflamed HSO infected by SARS-CoV-2 Reporter and SARS-CoV-2 Infection in Lungs of COVID-19 Patients. RNA extracted from the epidermal compartment of control HSO, HSO treated with the cytokine cocktail, and HSO infected with Reporter after pretreatment with the cytokine cocktail (TNF-α + IL-6 + IL-1β + IFN-γ; three HSO per group) was subjected to bulk RNA sequencing. (A) Increased transcriptomic signature in inflamed SHO after infection with SARS-CoV-2 Spike reporter. Differentially expressed genes (DEGs; FC > 2; P < 0.05; FDR <0.01) in HSO treated with cytokine cocktail compared to HSO ptretread with cytokine cocktail followed by virus infection ; Reactome pathway enrichment analysis (GSEA) of unique 596 upregulated DEGs induced by virus. Reactome pathway enrichment analysis (GSEA) of unique 596 DEGs, with highlighted Rho GTPase signaling pathways. (B). Shared upregulated DEGs in HSO ptretread with cytokine cocktail followed by virus infection and DEGs identified in lungs of deceased COVID-19 patients (top 500 upregulated DEGs, GSE150316). Reactome pathway enrichment analysis (GSEA) of overlapping 148 DEGs, with highlighted pathway related to SARS-CoV-2 infection.

Because fluorescent reporter (such as GFP or RFP) expression has been shown to have minimal effects on key cellular metabolic pathways and does not broadly alter host transcriptional programs [35-36], we assumed that transcriptomic changes observed in infected skin organoids primarily reflect downstream signaling induced by Spike protein binding to keratinocytes ACE2 in an inflamed HSO, rather than cell response to reporter DNA or expression of RFP.

RNA isolated from control HSO, cytokine-treated HSO, and cytokine-inflamed HSO infected with the SARS-CoV-2 reporter lentivirus was subjected to bulk RNA sequencing (GSE304119), and differentially expressed genes (DEGs, FC >2; P<0.01, FDR <0.05) were identified. As expected, the combination of inflammatory cytokines induced substantial transcriptomic changes in HSO (2,473 up- and downregulated DEGs). Notably, infection with the reporter lentivirus further broadened the cytokine-induced response, increasing the total number of DEGs to 3,383 (Fig. 5A). Gene set enrichment analysis (GSEA) of 596 uniquely upregulated DEGs induced by the reporter virus in inflamed HSO revealed strong enrichment of pathways related to inflammation, including innate immune responses, cytokine signaling, and neutrophil degranulation, as well as multiple pathways associated with the Rho (Ras homolog family) GTPase cycle (Fig. 5A, GSEA table). These are important and intersting findings as Rho GTPases, the key regulators of actin cytoskeleton dynamics and cellular architecture [37], play critical roles in viral infection including SARS-CoV-2 infection by facilitating viral entry, promoting replication, and regulating vesicular transport processes required for viral assembly and release [38-39]. Consistent with this, recent phosphoproteomic analyses of host cells infected with different SARS-CoV-2 variants has demonstrated the increased expression and activation of Rho GTPases as one of the major changes after infection [39].

Next, we compared the transcriptomes of lungs from deceased COVID-19 patients with the transcriptomic changes induced in cytokine-inflamed HSO following reporter lentivirus infection. DEGs lists in lung tissue of patients with COVID-19 were obtained from publicly available datasets curated within Enrichr, a comprehensive gene set enrichment analysis web server (https://maayanlab.cloud/Enrichr/) developed by Dr. Ma’ayan laboratory at Mount Sinai Center for Bioinformatics (Mount Sinai Center, New York) [40]. Comparison of available on Enrichr webserver top 500 upregulated DEGs from COVID-19-infected lung tissues with all 1671 upregulated DEGs from cytokine-treated, infected by Spike-pseudotyped virus skin organoids revealed a substantial shared transcriptional signature comprising 148 overlapping genes (Fig. 5B). GSEA of these common DEGs identified significant enrichment of pathways related to inflammation, innate immune responses, and adaptive immune activation. Importantly, the analysis also revealed the gene set associated with SARS-CoV-2–induced activation of immune pathways indicating some similarity between pulmonary COVID-19 signature and the signaling driven by Spike-pseudotyped lentiviral reporter in inflamed HSO models. Given that the lentiviral reporter system models viral entry but does not recapitulate full viral replication, these transcriptional changes likely reflect host responses to Spike–ACE2 interaction and downstream signaling. Overall, the transcriptome changes in HSO after infection by the SARS-CoV-2 reporter lentivirus further confirms the relevance of our models and supports its importance of studying skin responses to SARS-CoV-2 infection.

## 3. Discussion

This study provides experimental evidence that human skin, particularly under inflammatory conditions, can support SARS-CoV-2 attachment and entry. Using 3D human skin organoids (HSO) and neonatal skin explants, we show that epidermal keratinocytes express key viral entry factors, including ACE2 and TMPRSS2, and can be efficiently transduced by a Spike-pseudotyped reporter. Importantly, this susceptibility is markedly enhanced by proinflammatory cytokines.

A key strength of this work is the use of 3D skin models, which better recapitulate epidermal architecture and differentiation than conventional 2D systems [24],[32]. This is particularly relevant given conflicting reports on ACE2 and TMPRSS2 expression in keratinocytes. For example, Hertereau et al. [31] reported minimal ACE2 and no TMPRSS2 expression in 2D keratinocyte cultures. In contrast, we observed robust expression of both factors in 3D HSO and skin explants, consistent with studies demonstrating that epithelial differentiation state strongly influences ACE2 expression [9],[11],[41]. These findings underscore the importance of tissue context when assessing susceptibility to viral entry.

Our data further show that inflammatory signals reshape the molecular landscape required for viral entry. Cytokines associated with severe COVID-19 and chronic skin inflammation, including TNF-α, IL-1β, IL-6, and IFN-γ, significantly upregulated ACE2 and TMPRSS2, with strong synergistic effects when combined. Th1 cytokines (TNF-α + IL-17) preferentially induced ACE2, whereas Th2 cytokines (IL-4 + IL-13) increased TMPRSS2 expression. These findings are consistent with prior studies showing interferon-dependent induction of ACE2 [9],[31],[42-43] and cytokine-mediated upregulation of TMPRSS2 [43-44]. Functionally, this translated into increased Spike-mediated entry in inflamed skin models, particularly under Th1-skewed and cytokine storm–like conditions, while Th2 cytokines had minimal effects, consistent with their limited impact or even negative effect on ACE2 expression (our Fig. 3 and [45]). Together, these results suggest that inflamed skin, especially in Th1-dominant conditions such as psoriasis, may transiently acquire increased susceptibility to viral entry.

Transcriptomic analysis further supports the biological relevance of this model. Spike-mediated entry in inflamed HSO induced gene expression changes that partially overlapped with signatures observed in lung tissue from COVID-19 patients, particularly in pathways related to innate immunity and inflammation [46]. Although this overlap is incomplete, it suggests that Spike–ACE2 engagement in keratinocytes can initiate downstream signaling programs resembling aspects of host responses observed in vivo.

The role of skin as a potential entry site for SARS-CoV-2 remains incompletely understood. Previous studies have shown that keratinocytes can bind or internalize the virus, although infection is generally non-productive [31]. Our findings support a model in which inflamed or barrier-disrupted skin may permit viral attachment and entry without serving as a major site of viral replication. In addition, the enhanced inflammatory response observed following reporter infection suggests a potential feed-forward loop, where inflammation increases susceptibility to viral entry, which in turn amplifies inflammatory signaling, that may contribute to cutaneous manifestations such as vasculitis- or urticaria-like lesions in COVID-19 patients.

Several limitations should be considered. The Spike-pseudotyped lentiviral reporter that we used in our work, models viral entry but not the full viral life cycle. The neonatal skin and HSO have a less mature barrier than adult skin [47-48], which may overestimate susceptibility. In addition, SARS-CoV-2 entry is not solely dependent on ACE2 and TMPRSS2; alternative receptors, cofactors and proteases including cathepsin L (CTSL), neuropilin-1, heparan sulfate proteoglycans, all of which are expressed in keratinocytes as well as ASGR1, KREMEN1, and AXL receptor tyrosine kinase may contribute under specific conditions [49-50].

We also explored potential differences between African American and White non-Hispanic skin models. Unfortunately, relatively small sample size and inter-individual variability in ACE2 and TMPRSS2 expression limited our ability to detect variations in sensitivity of AA and WNH models to infection by SARS-CoV-2 reporter virus. Indeed, although we observed a trend toward stronger Th1-induced ACE2 expression in AA-derived HSOs, this did not translate into significant differences in viral entry. These findings should be interpreted cautiously and are consistent with evidence that disparities in COVID-19 outcomes are largely driven by social and structural factors rather than intrinsic biological differences [25-27].

In summary, our study demonstrates that inflammatory cues enhance the expression of key SARS-CoV-2 entry factors and increase susceptibility to Spike-mediated viral entry in human skin models. These findings identify inflammation as a critical modifier of epidermal permissiveness and suggest that the skin, particularly under inflammatory or barrier-compromised conditions, may represent a previously underappreciated interface in SARS-CoV-2 host interactions.

## 4 Materials and Methods

### Human Skin Explants

Neonatal foreskin samples provided by the SBDRC STEM Core were obtained from routine circumcisions performed at Prentice Women and Children’s Hospital in Chicago. Grouping into AA and WNH categories was based on mothers’ self-reported ethnicity. Published data indicate that self-reported AA ancestry correlates with genotyping at approximately 85%-90% [51].

### 3D Human Skin Organoids (HSO)

HSO provided by the SBDRC STEM Core were prepared using normal human epidermal keratinocytes (NHEKs) isolated from AA and WNH neonatal foreskins, as previously described [28]. Grouping was based on mothers’ self-reported ethnicity. HSO were cultured at the air–liquid interface for 10–12 days prior to treatment to promote keratinocyte differentiation and skin barrier establishment. In each experiment we used HSO generated from three-four independent primary keratinocyte lines/group.

### Treatment with Cytokines

Mature HSO with an established epidermis were treated for 7 days with individual cytokines: IL-1β, IL-6, IFN-γ, TNF-α (10 ng/mL each) or with their combination. In additional experiments, HSO were treated for 7 days with Th1 (IL-17 + TNF-α, 10 ng/mL each) or Th2 (IL-4 + IL-13, 30 ng/mL each) cytokine cocktails. Fresh cytokines were added every second day during media changes.

### Infection with SARS-CoV-2 Spike protein-pseudotyped Reporter Lentivirus

Standard lentiviral vectors are typically pseudotyped with vesicular stomatitis virus glycoprotein (VSV-G), which mediates viral entry through binding to widely expressed phospholipids rather than specific host receptors. To achieve ACE2-dependent cell entry, we generated a Tomato-expressing reporter lentivirus pseudotyped with the SARS-CoV-2 Spike protein by replacing VSV-G with the Spike protein from the Wuhan-Hu-1 strain as described [34].

HSO and skin explants were pretreated for four days with vehicle control (0.1% BSA in PBS) or with cytokine cocktails added to the culture medium: TNF-α + IL-1β + IL-6 + IFN-γ, Th1, or Th2 cytokines. Following pretreatment, HSO or skin explants were infected by topical application of the SARS-CoV-2 Spike-pseudotyped Tomato (Red fluorescent protein, RFP) reporter lentivirus. Samples were harvested three days post-infection, an optimal time point for detection of Tomato fluorescence, and were either immediately used for imaging and quantitative fluorescence analysis or flash-frozen for downstream biochemical analyses of Tomato expression at the mRNA level.

### Protein Extraction and Western blotting

Neonatal foreskin samples were homogenized using a ceramic bead homogenizer (PRO250) followed by centrifugation for 1 minute at 8000xg. HSO epidermis was homogenized by syringing. Proteins were extracted using Urea Sample Buffer (USB), consisting of consisting of 80% deionized Urea, 1% SDS, 10% glycerol, 0.06M Tris HCl (pH 6.8) and 0.001% Pyronin Y. Protein concentration was determined by Amido Black Assay on a VICTOR X plate reader (PerkinElmer, Shelton, CT, USA).

Proteins were resolved by SDS-PAGE and transferred to nitrocellulose membranes. The membranes were blocked by using 5% non-fat milk (Bio-Rad, Hercules, CA, USA) in Tris-Buffered Saline with Tween-20 (TBST) and incubated overnight at 4 °C with primary antibodies followed by secondary antibodies incubation for 1 hour at room temperature. All washes between antibodies were done in Phosphate Buffered Saline with Tween-20 (PBST) for 15 minutes 3x. Membranes were imaged using AZURE ECL reagents and the AZURE Chemidoc digital imager (AZURE Biosystems, Dublin, CA, USA). GAPDH antibodies were used to normalize for equal sample loading. Membranes were re-probed with GAPDH Ab after incubating in Stripping Buffer (Bio-Rad, Hercules, CA, USA) for 20 minutes, washing twice with TBST and blocking with 5% non-fat milk for 60 minutes. The differences in the protein expression between control and treated HSO were assessed by unpaired two-tailed Welch’s t-test.

We used following antibodies for Western blot analyses: ACE2 (Cell Sig #4355S, Cell Signaling, Danvers, MA, USA), TMPRSS2 (#sc515727, Santa Cruz, Dallas, TX, USA), GAPDH (Cell Sig #2118S, Cell Signaling, Danvers, MA, USA).

### RNA extraction and qRT-PCR

Skin samples were homogenized by PRO250 homogenizer (Pro Scientific, Oxford, CT, USA) on ice followed by syringing in QIAzol (Qiagen) buffer. HSO epidermis was homogenized by syringing in QIAzol buffer. RNA was extracted following the miRNeasy kit (Qiagen, Germantown, MD, USA) protocol. RNA concentrations were analyzed using a Nanodrop 2000 (Thermo-Fisher, Waltham, MA, USA) or RNA Qubit analysis (NUSeq Core, Northwestern University). cDNA synthesis was performed using random hexamers and M-MLV Reverse Transcriptase (Invitrogen, Carlsbad, CA, USA) according to the manufacturer’s protocol. The qRT-PCR for target genes was performed on a LightCycler 96 system (Roche Life Science, Penzberg, Germany) with SYBR Green I PCR Master Mix (Roche Life Science, Penzberg, Germany). House-keeping gene RPL27 was used for normalization, relative expression was calculated by the standard 2^(-ΔΔCt) approach [52]. Primers were designed using NCBI BLAST (http://www.ncbi.nlm.nih.gov/tools/primer-blast). The differences in the expression between control and treated AA and WNH samples were assessed by using unpaired two-tailed Welch’s t-test.

### RNA-sequencing and analysis of data

RNA-seq was performed by LC Sciences (Houston, TX, USA) on the Illumina Novaseq™ 6000 platform (paired-end reads, 150 bp). Per-base quality was assessed with FastQC software (http://www.bioinformatics.babraham.ac.uk/projects/fastqc/). Alignment of fastq files to the human genome (ftp://ftp.ensembl.org/pub/release-96/fasta/homosapiens/dna/) was done with HISAT2. The assembly of mapped reads of each sample and estimation of the expression levels was performed using StringTie. DEGs were selected at p-value ≤ 0.01 (FDR <0.05) and FC >2. GSEA analysis was performed online using

Molecular Signatures Database (MSigDB) v2026.1.Hs updated in January 2026, a joint project of UC San Diego and Broad Institute [53].

Lists of differentially expressed genes (DEGs) from lungs of patients with COVID-19 were obtained from publicly available datasets and queried using Enrichr (https://maayanlab.cloud/Enrichr/), a gene set analysis platform developed by Dr. Ma’ayan’s Laboratory at Mount Sinai Center for Bioinformatics, New York. These COVID-19 lung DEGs, provided as curated gene lists within Enrichr, were compared with RNA-seq data generated from HSO treated with cytokine storm-associated cytokines and infected with lentivirus pseudotyped with the SARS-CoV-2 spike protein.

### Multiphoton microscopy imaging of TD-Tomato infection in HSO and explant samples

Imaging was conducted using a Nikon A1R-MP microscope at the Northwestern Nikon Imaging Center. HSO samples were collected post-infection, mounted on charged slides, rinsed with PBST buffer, treated with 4′,6-diamidino-2-phenylindol (DAPI) for 10 minutes for nuclear staining and covered with glass coverslips in Gelvatol to prevent fluorescent signals fading. Neonatal skin explants were collected post-infection, permeabilized using 0.1% Triton X-100, stained for 20 minutes using DAPI and mounted to glass slides in Gelvatol. Image analysis was conducted using the associated Nikon Software and image processing software Image J (FIJI).

## Supplementary Materials

N/A

## Author Contributions

Conceptualization, I.B.; methodology, D.T., P.G., D.A.C., A.K., B.S., P.B., B.P.W., I.B.; software, D.T.; validation, D.T., P.G., D.A.C., A.K., B.S., P.B.; formal analysis, D.T., P.G., D.A.C., A.K., B.S., B.P.W., I.B.; investigation, D.T., P.G., D.A.C., A.K., I.B.; resources, I.B.; data curation, I.B. D.T; writing—original draft preparation, D.T., P.G., D.A.C., I.B.; writing—review and editing, D.T., P.G., D.A.C, I.B.; visualization, D.T., P.G., D.A.C., B.S., I.B; supervision, I.B.; project administration, I.B.; funding acquisition, I.B.. All authors have read and agreed to the published version of the manuscript.

## Institutional Review Board (IRB) Statement

The study was approved under IRB# STU00009443 by Northwestern University Institutional Review Board with a waiver of consent for the collection of neonatal foreskins and surgical discard skin tissues. All human skin tissues and human primary keratinocytes were provided by the SBDRC STEM Core through the IRB-approved Dermatology Tissue Acquisition and Repository services - a federally-compliant biorepository under IRB purview at Northwestern University.

## Informed Consent Statement

Informed consent for human biospecimens was obtained in writing following all local and federal regulations under STU00009443 approved by the Northwestern University Institutional Review Board, with a waiver of consent for the collection of neonatal foreskins and surgical discard skin tissues.

## Data Availability Statement

All raw and preprocessed RNA-Seq data used in this study were deposited in the GEO database with the GEO accession number GSE120783. All other data generated or analyzed during this study are included in this article.

## Acknowledgments

We acknowledge Northwestern University Skin Biology and Diseases Resource-Based Center (SBDRC) of the National Institutes of Health STEM and GET iN Cores for technical support. We are also thankful to Korvell Russell for technical assistance with skin organoids.

## Abbreviations

The following abbreviations are used in this manuscript:

AA: African American
ACE2: angiotensin converting enzyme 2
AD: atopic dermatitis
BSA: bovine serum albumin
COVID-19: coronavirus disease 2019
CTSL: Cathepsin L
DAPI: 4′,6-diamidino-2-phenylindole
IFNγ: interferon gamma
IL: Interleukin
HSO: human skin organoid
NHEK: normal human epidermal keratinocytes
PBST: Phosphate-buffered Saline with Tween-20
RFP: red fluorescent protein
SARS-CoV-2: severe acute respiratory syndrome coronavirus 2
SDS-PAGE: Sodium Dodecyl Sulfate–Polyacrylamide Gel Electrophoresis
TBST: Tris-Buffered Saline with Tween-20
TMPRSS2: Transmembrane serine protease 2
TU: Transduction units
USB: Urea Sample Buffer
VSV-G: vesicular stomatitis virus glycoprotein
WNH: White non-Hispanic

## Disclaimer/Publisher’s Note

The statements, opinions and data contained in all publications are solely those of the individual author(s) and contributor(s) and not of MDPI and/or the editor(s). MDPI and/or the editor(s) disclaim responsibility for any injury to people or property resulting from any ideas, methods, instructions or products referred to in the content.

## References

1. Hoffmann M., Kleine-Weber H., Pöhlmann S. A multibasic cleavage site in the spike protein of SARS-CoV-2 is essential for infection of human lung cells. Molecular cell. 2020; 78(4):779–84. doi: 10.1016/j.molcel.2020.04.022

2. Jackson C.B., Farzan M., Chen B., Choe H. Mechanisms of SARS-CoV-2 entry into cells. Nature reviews Molecular cell biology. 2022; 23(1):3–20. doi: 10.1038/s41580-021-00418-x

3. Barthe M., Hertereau L., Lamghari N., Osman-Ponchet H., Braud V.M. Receptors and cofactors that contribute to SARS-CoV-2 entry: can skin be an alternative route of entry? International journal of molecular sciences. 2023; 24(7):6253. doi: 10.3390/ijms24076253

4. Bittmann S., Weissenstein A., Luchter E., Villalon G., Moschüring-Alieva E. ACE-2-Receptors of the Epidermis, Dermal Vascular Walls and Sebaceous Gland Cells: The Way of COVID-19 Entry into the Body. J Regen Biol Med. 2020; 2(3):1–3. doi: 10.37191/Mapsci-2582-385X-2(3)-029

5. Kaplan N., Gonzalez E., Peng H., Batlle D., Lavker R.M. Emerging importance of ACE2 in external stratified epithelial tissues. Mol Cell Endocrinol. 2021; 529, 111260. doi: 10.1016/j.mce.2021.111260

6. Ma D., Chen C.B., Jhanji V., Xu C., Yuan X.L., Liang J.J., Huang Y., Cen L.P., Ng T.K. Expression of SARS-CoV-2 receptor ACE2 and TMPRSS2 in human primary conjunctival and pterygium cell lines and in mouse cornea. Eye (Lond). 2020; 34(7):1212–1219. doi: 10.1038/s41433-020-0939-4.

7. Radzikowska U., Ding M., Tan G., Zhakparov D., Peng Y., Wawrzyniak P., Wang M., Li S., Morita H., Altunbulakli C., Reiger M., Neumann A.U., Lunjani N., Traidl-Hoffmann C., Nadeau K.C., O’Mahony L., Akdis C., Sokolowska M. Distribution of ACE2, CD147, CD26, and other SARS-CoV-2 associated molecules in tissues and immune cells in health and in asthma, COPD, obesity, hypertension, and COVID-19 risk factors. Allergy. 2020; 75(11), 2829–2845. doi: 10.1111/all.14429

8. Xue X., Mi Z., Wang Z., Pang Z., Liu H., Zhang F. High Expression of ACE2 on Keratinocytes Reveals Skin as a Potential Target for SARS-CoV-2. J Invest Dermatol. 2021; 141(1), 206–209 e201. doi: 10.1016/j.jid.2020.05.087

9. Lin L., Zeng F., Mai L., Gao M., Fang Z., Wu B., Huang S., Shi H., He J., Liu Y., Li X., Li Z., Han Y., Yan Z. Expression of ACE2, TMPRSS2, and SARS-CoV-2 nucleocapsid protein in gastrointestinal tissues from COVID-19 patients and association with gastrointestinal symptoms. Am J Med Sci. 2023 Dec;366(6):430–437. doi: 10.1016/j.amjms.2023.08.014

10. Konturek P.C., Harsch I.A., Neurath M.F., Zopf Y. COVID-19 - more than respiratory disease: a gastroenterologist’s perspective. J Physiol Pharmacol. 2020; 71(2). doi: 10.26402/jpp.2020.2.02.

11. Sun Y., Zhou R., Zhang H., Rong L., Zhou W., Liang Y., Li Q. Skin is a potential host of SARS-CoV-2: A clinical, single-cell transcriptome-profiling and histologic study. J Am Acad Dermatol. 2020; 83(6), 1755–1757. doi: 10.1016/j.jaad.2020.08.057

12. Lili L.N., Klopot A., Readhead B., Baida G., Dudley J.T., Budunova, I. Transcriptomic Network Interactions in Human Skin Treated with Topical Glucocorticoid Clobetasol Propionate. J Invest Dermatol. 2019; 139(11), 2281–2291. doi: 10.1016/j.jid.2019.04.021

13. Cazzato G., Cascardi E., Colagrande A., Foti C., Stellacci A., Marrone M., Ingravallo G., Arezzo F., Loizzi V., Solimando A.G., et al. SARS-CoV-2 and Skin: New Insights and Perspectives. Biomolecules. 2022; 12(9):1212. doi: 10.3390/biom12091212.

14. Brahimi N., Croitoru D., Saidoune F., Zabihi H., Gilliet M., Piguet V. From Viral Infection to Skin Affliction: Unveiling Mechanisms of Cutaneous Manifestations in COVID-19 and Post-COVID Conditions. J Invest Dermatol. 2025; 145(2), 257–265. doi: 10.1016/j.jid.2024.03.047

15. Novak N., Peng W., Naegeli M.C., Galvan C., Kolm-Djamei I., Bruggen C., Cabanillas B., Schmid-Grendelmeier P., Catala, A. SARS-CoV-2, COVID-19, skin and immunology - What do we know so far? Allergy. 2021; 76(3), 698–713. doi: 10.1111/all.14498

16. Visconti A., Bataille V., Rossi N., Kluk J., Murphy R., Puig S., Nambi R., Bowyer R.C.E., Murray B., Bournot A., et al. Diagnostic value of cutaneous manifestation of SARS-CoV-2 infection. Br J Dermatol. 2021; 184(5), 880–887. doi: 10.1111/bjd.19807

17. Bian X.W., COVID-19 Pathology Team. Autopsy of COVID-19 patients in China. Natl Sci Rev. 2020; 7(9), 1414–1418. doi: 10.1093/nsr/nwaa123

18. Jamiolkowski D., Muhleisen B., Muller S., Navarini A.A., Tzankov A., Roider, E. SARS-CoV-2 PCR testing of skin for COVID-19 diagnostics: a case report. Lancet. 2020; 396(10251), 598–599. doi: 10.1016/S0140-6736(20)31754-2

19. Cho Y.A., Han H., Won S., Lim J.W., Sung J.Y., Kim C.Y., Yu D.A., Lee Y.W., Choe Y.B. Covid-19 susceptibility, severity, and vaccine effectiveness in patients with psoriasis: a nationwide cohort study in south Korea. Scientific Reports. 2025; 15(1):25608. doi: 10.1038/s41598-025-06495-8

20. Krueger J.G., Murrell D.F., Garcet S., Navrazhina K., Lee P.C., Muscianisi E., Blauvelt A. Secukinumab lowers expression of ACE2 in affected skin of patients with psoriasis. J Allergy Clin Immunol. 2021; 147(3), 1107–1109 e1102. doi: 10.1016/j.jaci.2020.09.021

21. Patrick M.T., Zhang H., Wasikowski R., Prens E.P., Weidinger S., Gudjonsson J.E., Elder J.T., He K., Tsoi L.C. Associations between COVID-19 and skin conditions identified through epidemiology and genomic studies. J Allergy Clin Immunol. 2021; 147(3), 857–869 e857. doi: 10.1016/j.jaci.2021.01.006

22. Singh M.K., Mobeen A., Chandra A., Joshi S., Ramachandran S. A meta-analysis of comorbidities in COVID-19: Which diseases increase the susceptibility of SARS-CoV-2 infection? Comput Biol Med. 2021; 130, 104219. doi: 10.1016/j.compbiomed.2021.104219

23. Tembhre M.K., Parihar A.S., Sharma V.K., Imran S., Bhari N., Lakshmy R., Bhalla A. Enhanced expression of angiotensin-converting enzyme 2 in psoriatic skin and its upregulation in keratinocytes by interferon-gamma: implication of inflammatory milieu in skin tropism of SARS-CoV-2. Br J Dermatol. 2021; 184(3), 577–579. doi: 10.1111/bjd.19670

24. Eyerich K., Brown S.J., Perez White B.E., Tanaka R.J., Bissonette R., Dhar S., Bieber T., Hijnen D.J., Guttman-Yassky E., Irvine A., Thyssen J.P., Vestergaard C., Werfel T., Wollenberg A., Paller A.S., Reynolds N.J. Human and computational models of atopic dermatitis: A review and perspectives by an expert panel of the International Eczema Council. J Allergy Clin Immunol. 2019; 143(1), 36–45. doi: 10.1016/j.jaci.2018.10.033

25. Magesh S., John D., Li W.T., Li Y., Mattingly-App A., Jain S., Chang E.Y., Ongkeko W.M. Disparities in COVID-19 Outcomes by Race, Ethnicity, and Socioeconomic Status: A Systematic-Review and Meta-analysis. JAMA Netw Open. 2021; 4(11), e2134147. doi: 10.1001/jamanetworkopen.2021.34147

26. Shortreed S.M., Gray R., Akosile M.A., Walker R.L., Fuller S., Temposky L., Fortmann S.P., Albertson-Junkans L., Floyd J.S., Bayliss E.A., Harrington L.B., Lee M.H., Dublin S. Increased COVID-19 Infection Risk Drives Racial and Ethnic Disparities in Severe COVID-19 Outcomes. J Racial Ethn Health Disparities. 2023; 10(1), 149–159. doi: 10.1007/s40615-021-01205-2

27. Zelner J., Trangucci R., Naraharisetti R., Cao A., Malosh R., Broen K., Masters N., Delamater P. Racial Disparities in Coronavirus Disease 2019 (COVID-19) Mortality Are Driven by Unequal Infection Risks. Clin Infect Dis. 2021; 72(5), e88–e95. doi: 10.1093/cid/ciaa1723

28. Klopot A., Baida G., Kel A., Tsoi L.C., Perez White B.E., Budunova I. Transcriptome Analysis Reveals Intrinsic Proinflammatory Signaling in Healthy African American Skin. J Invest Dermatol. 2022; 142(5), 1360–1371 e1315. doi: 10.1016/j.jid.2021.09.031

29. Dharra R., Kumar Sharma A., Datta S. Emerging aspects of cytokine storm in COVID-19: The role of proinflammatory cytokines and therapeutic prospects. Cytokine. 2023; 169, 156287. doi: 10.1016/j.cyto.2023.156287

30. Paranga T.G., Mitu I., Pavel-Tanasa M., Rosu M.F., Miftode I.L., Constantinescu D., Obreja M., Plesca C.E., Miftode E. Cytokine Storm in COVID-19: Exploring IL-6 Signaling and Cytokine-Microbiome Interactions as Emerging Therapeutic Approaches. Int J Mol Sci. 2024; 25(21). doi: 10.3390/ijms252111411

31. Hertereau L., Barthe M., Lamghari N., Merida P., Pommier G., Pinel E., Swain J., Muriaux D., Osman-Ponchet H., Braud V.M. Human keratinocytes exhibit limited potential for SARS-CoV-2 infection despite ACE2 and mature cathepsin L expression. JID Innovations. 2025; 6(2):100447. doi: 10.1016/j.xjidi.2025.100447

32. Desmet E., Ramadhas A., Lambert J., Van Gele M. In vitro psoriasis models with focus on reconstructed skin models as promising tools in psoriasis research. Experimental biology and medicine. 2017; 242(11):1158–69. doi: 10.1177/1535370217710637

33. Criado P.R., Ianhez M., Silva de Castro C.C., Talhari C., Ramos P.M., Miot H.A. COVID-19 and skin diseases: results from a survey of 843 patients with atopic dermatitis, psoriasis, vitiligo and chronic urticaria. J Eur Acad Dermatol Venereol. 2022; 36(1):e1–e3. doi: 10.1111/jdv.17635

34. Tandon R., Mitra D., Sharma P., McCandless M.G., Stray S.J., Bates J.T., Marshall G.D. Effective screening of SARS-CoV-2 neutralizing antibodies in patient serum using lentivirus particles pseudotyped with SARS-CoV-2 spike glycoprotein. Sci Rep. 2020; 10(1), 19076. doi: 10.1038/s41598-020-76135-w

35. Soboleski M.R., Oaks J., Halford W.P. Green fluorescent protein is a quantitative reporter of gene expression in individual eukaryotic cells. FASEB J. 2005; 19(3), 440–442. doi: 10.1096/fj.04-3180fje

36. Yang J., Wang N., Chen D., Yu J., Pan Q., Wang D., Liu J., Shi X., Dong X., Cao H., Li L., Li L. The Impact of GFP Reporter Gene Transduction and Expression on Metabolomics of Placental Mesenchymal Stem Cells Determined by UHPLC-Q/TOF-MS. Stem Cells Int. 2017; 3167985. doi: 10.1155/2017/3167985

37. Olson M.F. Rho GTPases, their post-translational modifications, disease-associated mutations and pharmacological inhibitors. Small GTPases. 2018; 9(3):203–215. doi: 10.1080/21541248.2016.1218407

38. Zhang B., Li S., Ding J., Guo J., Ma Z., Duan H. Rho-GTPases subfamily: cellular defectors orchestrating viral infection. Cell Mol Biol Lett. 2025; 30(1):55. doi: 10.1186/s11658-025-00722-w.

39. Huuskonen S., Liu X., Pöhner I., Redchuk T., Salokas K., Lundberg R., Maljanen S., Belik M., Reinholm A., Kolehmainen P., et al. The comprehensive SARS-CoV-2 ‘hijackome’ knowledge base. Cell Discov. 2024; 10(1):125. doi: 10.1038/s41421-024-00748-y.

40. Kuleshov M.V., Jones M.R., Rouillard A.D., Fernandez N.F., Duan Q., Wang Z., Koplev S., Jenkins S.L., Jagodnik K.M., Lachmann A., et al. Enrichr: a comprehensive gene set enrichment analysis web server 2016 update. Nucleic acids research. 2016; 44(W1):W90–7. doi: 10.1093/nar/gkw377

41. Sungnak W., Huang N., Bécavin C., Berg M., Queen R., Litvinukova M., Talavera-López C., Maatz H., Reichart D., Sampaziotis F., Worlock K.B. SARS-CoV-2 entry factors are highly expressed in nasal epithelial cells together with innate immune genes. Nature medicine. 2020; 26(5):681–7. doi: 10.1038/s41591-020-0868-6

42. Saheb Sharif-Askari F., Goel S., Saheb Sharif-Askari N., Hafezi S., Al Heialy S., Hachim M.Y., Hachim I.Y., Mahboub B., Salameh L., Abdelrazig M., et al. Asthma Associated Cytokines Regulate the Expression of SARS-CoV-2 Receptor ACE2 in the Lung Tissue of Asthmatic Patients. Front Immunol. 2022; 12:796094. doi: 10.3389/fimmu.2021.796094

43. Sajuthi S.P., DeFord P., Li Y., Jackson N.D., Montgomery M.T., Everman J.L., Rios C.L., Pruesse E., Nolin J.D., Plender E.G., Wechsler M.E., et al. Type 2 and interferon inflammation regulate SARS-CoV-2 entry factor expression in the airway epithelium. Nat Commun. 2020; 11(1):5139. doi: 10.1038/s41467-020-18781-2

44. Cioccarelli C., Sánchez-Rodríguez R., Angioni R., Venegas F.C., Bertoldi N., Munari F., Cattelan A., Molon B., Viola A. IL1β Promotes TMPRSS2 Expression and SARS-CoV-2 Cell Entry Through the p38 MAPK-GATA2 Axis. Front Immunol. 2021; 12:781352. doi: 10.3389/fimmu.2021.781352

45. Kimura H., Francisco D., Conway M., Martinez F.D., Vercelli D., Polverino F., Billheimer D., Kraft M. Type 2 inflammation modulates ACE2 and TMPRSS2 in airway epithelial cells. J Allergy Clin Immunol. 2020; 146(1):80–88.e8. doi: 10.1016/j.jaci.2020.05.004

46. Gordon D.E., Jang G.M., Bouhaddou M., Xu J., Obernier K., White K.M., O’Meara M.J., Rezelj V.V., Guo J.Z., Swaney D.L., et al. A SARS-CoV-2 protein interaction map reveals targets for drug repurposing. Nature. 2020; 583(7816):459–468. doi: 10.1038/s41586-020-2286-9

47. Visscher M.O., Carr A.N., Narendran V. Epidermal Immunity and Function: Origin in Neonatal Skin. Front Mol Biosci. 2022; 9, 894496. doi: 10.3389/fmolb.2022.894496

48. Visscher M.O., Carr A.N., Winget J., Huggins T., Bascom C.C., Isfort R., Lammers K., Narendran V. Biomarkers of neonatal skin barrier adaptation reveal substantial differences compared to adult skin. Pediatr Res. 2021; 89(5):1208–1215. doi: 10.1038/s41390-020-1035-y.

49. Cantuti-Castelvetri L., Ojha R., Pedro L.D., Djannatian M., Franz J., Kuivanen S., van der Meer F., Kallio K., Kaya T., Anastasina M., et al. Neuropilin-1 facilitates SARS-CoV-2 cell entry and infectivity. Science. 2020; 370(6518):856–860. doi: 10.1126/science.abd2985

50. Daly J.L., Simonetti B., Klein K., Chen K.E., Williamson M.K., Anton-Plagaro C., Shoemark D.K., Simon-Gracia L., Bauer M., Hollandi R., et al. Neuropilin-1 is a host factor for SARS-CoV-2 infection. Science, 2020; 370(6518), 861–865. doi: 10.1126/science.abd3072

51. Yaeger R., Avila-Bront A., Abdul K., Nolan P.C., Grann V.R., Birchette M.G., Choudhry S., Burchard E.G., Beckman K.B., Gorroochurn P., et al. Comparing genetic ancestry and self-described race in african americans born in the United States and in Africa. Cancer Epidemiol Biomarkers Prev. 2008; 17(6), 1329–1338. doi: 10.1158/1055-9965.EPI-07-2505

52. Livak K.J., Schmittgen T.D. Analysis of relative gene expression data using real-time quantitative PCR and the 2(-Delta Delta C(T)) Method. Methods. 2001; 25(4):402–8. doi: 10.1006/meth.2001.1262

53. Subramanian A., Tamayo P., Mootha V.K., Mukherjee S., Ebert B.L., Gillette M.A., Paulovich A., Pomeroy S.L., Golub T.R., Lander E.S., et al. Gene set enrichment analysis: a knowledge-based approach for interpreting genome-wide expression profiles. Proc Natl Acad Sci U S A. 2005; 102(43):15545–50. doi: 10.1073/pnas.0506580102

